# Modelling Large, Dynamic, and Heterogeneous Populations Using DNA Libraries

**DOI:** 10.1101/276873

**Authors:** Helena Andrade, Alvin K. Thomas, Weilin Lin, Francesco V. Reddavide, Yixin Zhang

## Abstract

The study of any population of large size and high diversity is limited by the lack of data and associated insights. For a pool of individuals, each associated with a unique characteristic feature, as the pool size grows, the possible interactions increase exponentially, quickly beyond the scope of computation, not to mention experimental manipulation and analysis. Herein, we report a facile RT-PCR-based method, to correlate the amplification curves with various DNA libraries of defined diversity, and perform operations with groups of quaternary numbers as input and diversity as output. An attractive feature of this approach is the possibility of realizing parallel computation with an eventually unlimited number of variables. We demonstrate that DNA libraries can be used to model heterogeneous populations, exhibiting functions such as self-protection, subjected to biased expansion, and to evolve into complex structures. Moreover, the method can be applied to drug discovery using DNA-encoded chemical library (DECL) technology, to optimize selection conditions for identifying potent and specific bio-molecular interactions.

## INTRODUCTION

“*God has put a secret art into the force of Nature so as to enable it to fashion itself out of chaos into a perfect world system”* Immanuel Kant

Large and dynamic populations of high diversity are difficult to sample, analyze, and model^1,2^. We propose that a synthetic DNA library represents the ideal medium for establishing experimental systems for modeling large populations of high complexity and dynamics. DNA libraries can be synthesized with controlled varying diversity, manipulated enzymatically, subjected to growth using polymerase chain reaction (PCR), and analyzed with various methods including real-time PCR (RT-PCR or qPCR)^3,4^, and sequencing^5^. For example, a randomized 20-base sequence (N20) contains more than one trillion different DNA sequences (420). Only 10 μL of 1 nM N20 solution has the size of the human population, and statistically each molecule represents a unique individual.

The design of DNA libraries to model large populations of various diversities is shown in **Fig. 1a**. 20-base sequences **X**_**n**_ (**n** is the length of degenerate sequence and reflects the library diversity) are flanked by primers **A** and **B**. We first carried out a simulation of the PCR amplification process of libraries of different diversities (**Fig. 1b** and **1c**). In a high diversity library, two fully complimentary sequences have an extremely low probability to encounter each other, thus generating probabilistically self-assembled mismatching pairs. We assume that during the PCR process, an **A-X-B** sequence has the same probability to assemble with any **A’- X’-B’** as well as with the primer **B’**, while each **A’-X’-B’** has the same probability to form a duplex with either **A-X-B** or **A**.

**Figure 1.**
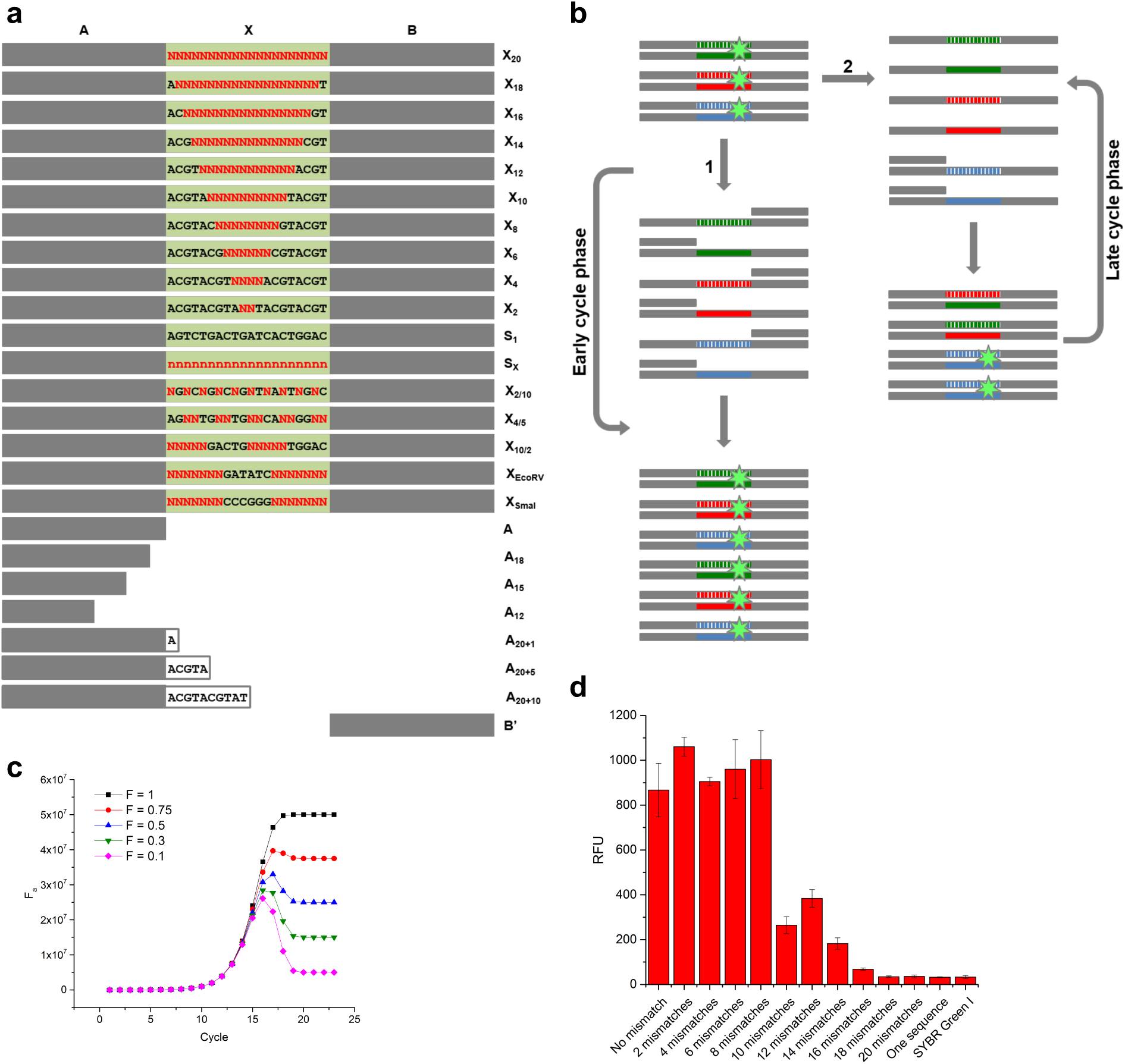
Amplification of DNA libraries of different diversities. (**a**) Design of DNA libraries and primers (**Extended Data Table 1**). (**b**) The formation of fully complimentary duplex in the initial PCR cycles (**1**) and mismatched duplex in the late PCR cycles (**2**). The mismatched duplex possesses lower F factor and lower affinity to fluorescence dye (green star) at high temperature (e.g., 72 °C of elongation temperature). (**c**) Simulation of PCR process using DNA samples of different diversity. (**d**) Mismatch effect on the fluorescence signal of 100 nM annealed products at 72 °C of elongation temperature.

We assigned a fully complimentary duplex with a functional factor (F) of 1.0. Mismatching leads to lower F values, e.g., **A-*X*_*10*_-B/A’-*X10’*-B’** and **A-*X*_*20*_-B/A’-*X20’*-B’** duplexes have F values of 0.5 and 0.1, respectively (the italic ***X*** and ***X’*** indicate mismatching in the randomized region). Interestingly, throughout the PCR cycles, although the production of full length DNA was not affected by using either **A-X_S1_-B** or various libraries as templates, the time courses of F values differed from each other dramatically. In the initial phase (**Fig. 1b path-1**), primer concentrations are much higher than those of full length DNA, probabilistically self-assembled **A-X-B/B’** And **A’-X’-B’/A** Are more abundant than **A-*X*-B/A’-*X’*-B’**. **A-X-B/B’** and **A’-X’-B’/A** produce **A-<u>X</u>-B/A’-<u>X’</u>-B’** in the presence of DNA polymerase (**<u>X</u>**<u>/</u>**<u>X’</u>** means the fully complimentary pair synthesized by DNA polymerase, according to the template). The newly synthesized fully complimentary **A-<u>X</u>-B/A’-<u>X</u>’-B’** possesses an F value of 1.0. In later phases (**Fig. 1b path-2**), the primers are consumed and full length DNAs are abundant. Hence, **A-X-B/B’** and **A’-X’-B’/A** are much less prevalent than **A-*X*-B/A’-*X*’-B’**. The duplexes generated by annealing form mismatching pairs with lower F values, in contrast to the duplexes produced by polymerase, where F =1.0. There is a turning point in the course of a high diversity library (**Fig. 1c**), as the increase of F caused by newly synthesized fully complementary duplexes equals the decrease of F caused by the re-annealing of DNA, which generates mismatched pairs through re-shuffling among the library members.

## RESULTS and Discussion

### Diversity Analysis

To relate F to an experimentally measurable parameter, we took advantage of the fact that when **A-X-B** and **A’-X’-B’** anneal quickly, the resulting duplexes with more mismatches are less stable at elongation temperature, thus leading to weaker dye binding and thus lower fluorescence. The effect of changing DNA population through SELEX cycles on RT-PCR amplification and melting curves has been used to monitor the selection process^6,7^. If the amplification curves can be correlated with various mixtures of sequences of defined diversities (**Fig. 1a**), we can perform computations with groups of quaternary numbers as input and diversity as output. An attractive feature of this approach is the possibility of realizing parallel computation of an eventually unlimited number of variables. We first analyzed the effect of mismatching on the binding of fluorescence dye. A remarkably diminished fluorescence signal was detected when the mismatching number is ≥ 10, a further decrease when that is ≥ 14 (**Fig. 1d** and **Extended Data Fig. 1**). Therefore, with the same amount of DNA duplexes, the fluorescence intensity reflects sequence mismatching, within the course monitored by RT-PCR. It is important to note that a standard RT-PCR protocol uses high primer concentrations to ensure the robustness of the assay. In our experiments, we adjusted primer concentrations to the range where the primer concentrations exhibited a linear correlation with the final signal intensity (**Extended Data Fig. 2**).

**Figure 2.**
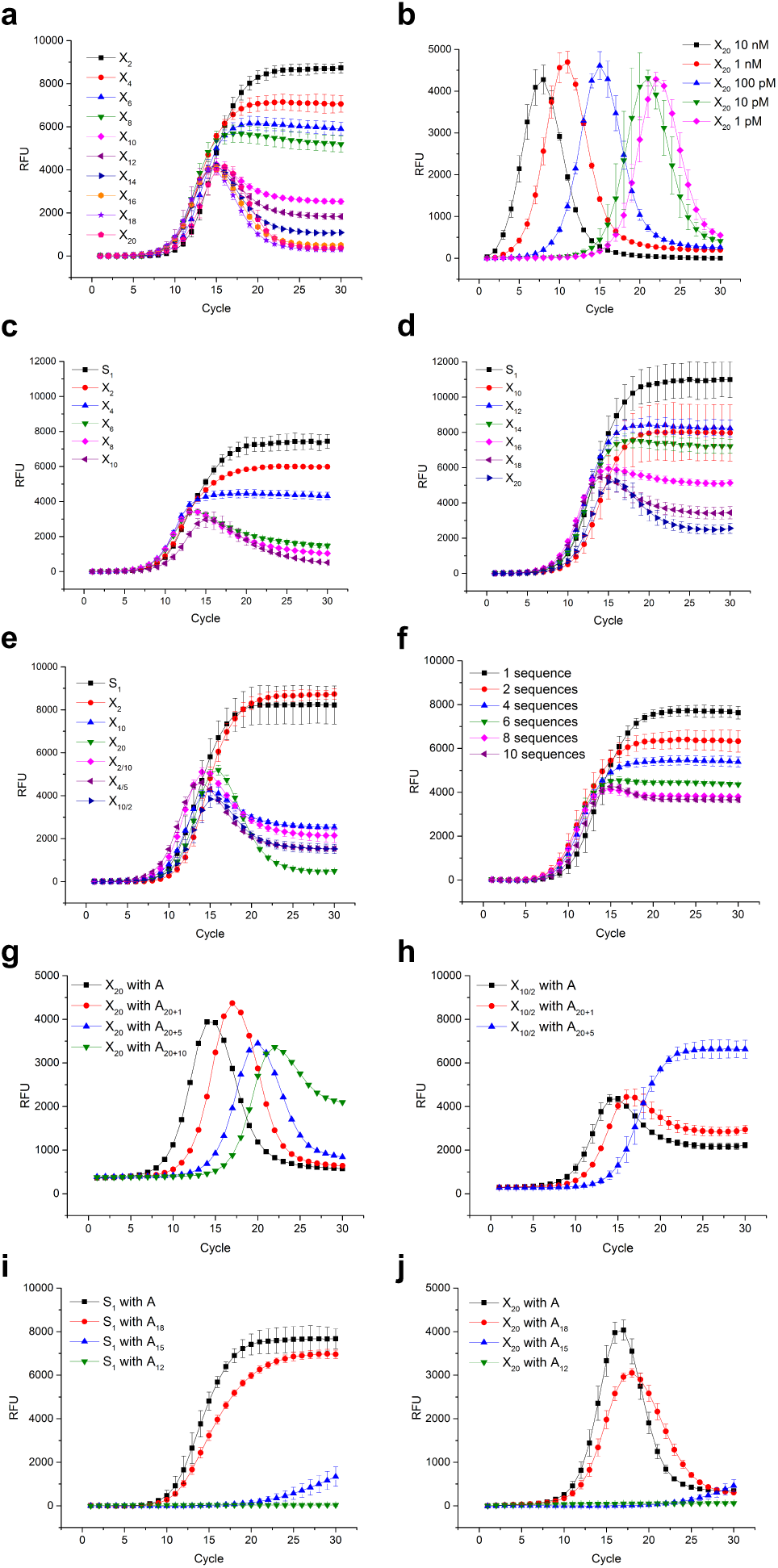
Diversity analysis and biased population growth. (**a**) RT-PCR experiments using DNA of different diversities (from two to 20 random positions) as templates. (**b**) RT-PCR experiments using different concentrations of **A-X_20_-B** as template. RT-PCR amplification with elongation at 74 (**c**) and at 68 °C (**d**). (**e**) RT-PCR experiments using different libraries of same diversity. **X**_**10**_, **X**_**10/2**_, **X**_**2/10**_, and **X**_**4/5**_ possess the same diversity but different distribution of partially degenerated segm**a**ents in the sequence. (**f**) RT-PCR experiments using a small number of different sequences. The sequences have been designed to possess large difference (>15) between any pair. Manipulation of the population growth using **A**, **A**_**20+1**_, **A**_**20+5**_, or **A**_**20+10**_ as primer with the libraries **X**_**20**_ (**g**) or **X**_**10/2**_ (**h**).

As shown in **Fig. 2a**, DNA libraries of different diversities have shown time courses resembling the simulated curves. The low diversity library **A-X_2_-B** contains only 16 different species and showed a RT-PCR profile like the sample containing only one sequence. RT-PCR of the highest diversity library, **A-X_20_-B,** exhibited a peak-shape curve. With increasing diversity, standard RT-PCR curves gradually transformed to peak-shaped curves. Interestingly, although the curves of high diversity libraries are remarkably different from those in classical RT-PCR measurements, the shifts of curves were correlated with the template concentration (**Fig. 2b** and **Extended Data Fig. 3**). Therefore, RT-PCR can be used to determine the sample concentration as well as to illustrate sample diversity.

**Figure 3.**
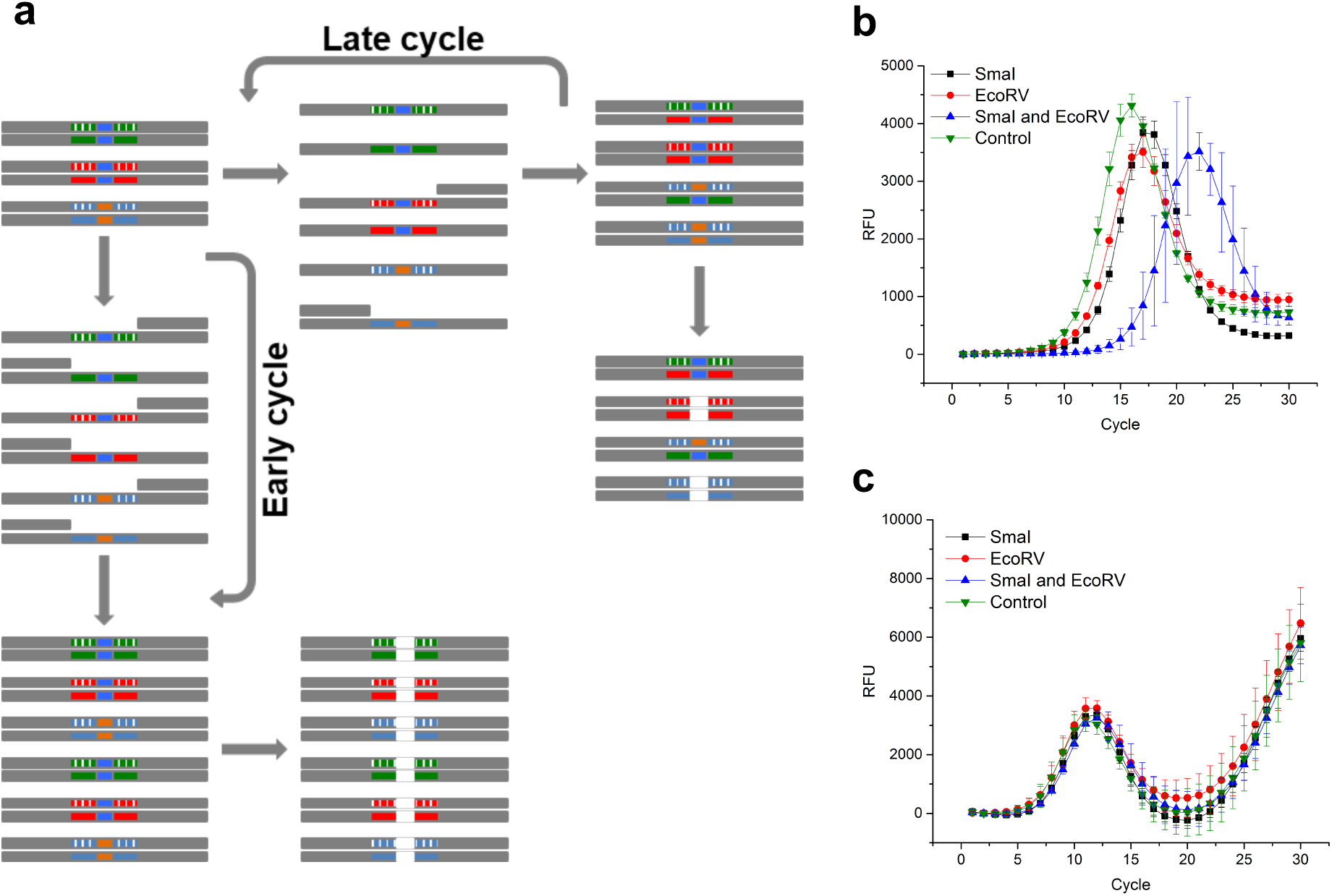
Diversity as a population protection mechanism. (**a**) Diversity of libraries with restriction sites has protective effect against endonuclease action. The fully complimentary duplex synthesized in the initial PCR cycles is more sensitive to endonucleases than the mismatched duplex generated in the late PCR cycles. The mixture of **A-XEcoRV-B** and **A-XSmal-B** was pre-treated with five (**b**) and 25 (**c**) PCR cycles. A small aliquot of PCR product was treated with EcoRV or Smal or both and the products were subjected to RT-PCR experiments.

The difference in diversity among populations does not necessarily fall into a wide range. The difference of RT-PCR curve among **X**_**4**_, **X**_**6**_ and **X**_**8**_ or among **X**_**10**_, **X**_**12**_ and **X**_**14**_ is relatively small, while **X**_**16**_, **X**_**18**_ and **X**_**20**_ cannot be distinguished from each other. The difference can be augmented through tuning the elongation temperature. The difference among **X**_**2**_, **X**_**4**_, **X**_**6**_, and **X**_**8**_ has been drastically increased at 74 °C, while **X**_**20**_, **X**_**18**_, **X**_**16**_, and **X**_**14**_ can be clearly distinguished from each other at 68 °C (**Fig. 2c** and **2d**). Therefore, besides covering a wide range of diversity, the condition can also be tuned to focus on a relatively narrow range.

We then tested different libraries of same diversity (**Fig. 1a** and **2e**). Although the partially degenerated part(s) are positioned very differently in the four libraries (**X**_**10**_, **X**_**2/10**_, **X**_**4/5**_, **X**_**10/2**_), the RT-PCR experiments resulted in similar profiles. For library **X**_**n**_, when n >16, each sequence in a 10 μL 1 nM sample (e.g., **X**_**18**_ and **X**_**20**_) is statistically unique, whereas each sequence in the **X**_**2/10**_ and **X**_**4/5**_ samples is represented a few thousand times. Interestingly, while **X**_**20**_, **X**_**18**_, **X**_**16**_, **X**_**2/10**_, and **X**_**4/5**_ possess randomized regions of similar length (18 to 20), the high diversity libraries (**X**_**20**_ and **X**_**18**_) produced curves clearly distinct from the medium diversity libraries (**X**_**2/10**_ and **X**_**4/5**_).

The difference between any two sequences in the **X**_**20**_ library can be of any number between zero and 20. We then designed a library of another type of diversity, a library containing *n* sub-libraries. In this library, all members in one sub-library are identical, but the difference between sub-libraries is very high. We generated library **Sx-10** of only ten sequences (**Extended Data Table 1**), which were designed to ensure the difference between any two sequences >15. If dynamic DNA duplex re-shuffling does not take place through the annealing process of each PCR cycle, the RT-PCR of **Sx-10** shall produce a curve resembling that of **X**_**20**_. Interestingly, the resulting curve (**Fig. 2f** and **Extended Data Fig. 4**) indicates that thermodynamic re-equilibration does play a role in this process, although it is far too inefficient to cause perfect matching among all sequences. Reducing the sub-library number transformed the curve to a standard RT-PCR curve gradually.

**Figure 4.**
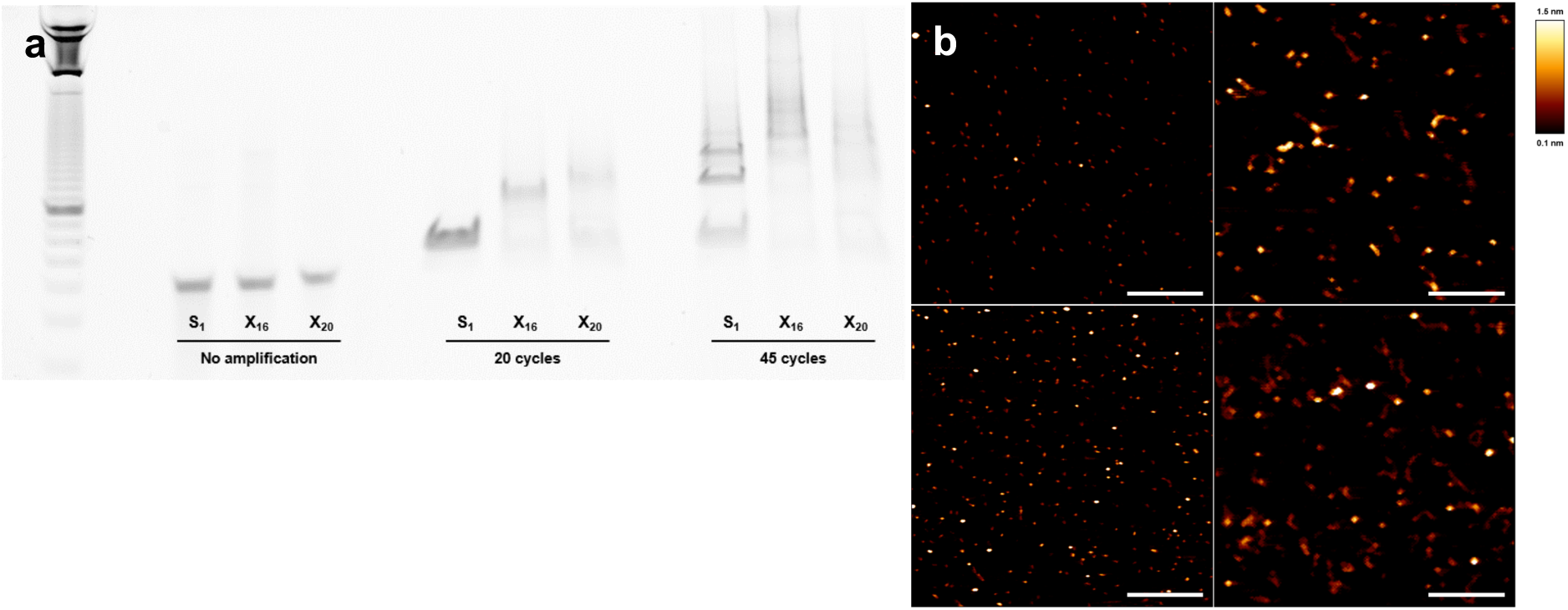
High-entropy structures formation. (**a**) DNA samples of different diversity (**S1**, **X**_**16**_, **X**_**20**_) were subjected to 20 or 45 PCR cycles and separated under denaturing conditions. (**b**) AFM analyses of PCR products of **S1** after 20 (upper-left) or 45 cycles (upper-right), **X**_**16**_ after 45 cycles (lower-left), and **X**_**20**_ after 45 cycles of amplification (lower-right). Images are 1 μM × 1 μM and the scale bar is 250 nm.

*Biased Population Growth*

We tested to reduce the diversity of DNA library through designing selective primers, while the biased amplification process could also be monitored using this RT-PCR method. **A**_**20+1**_, **A**_**20+5**_, and **A**_**20+10**_ (**Fig. 1**) should selectively anneal to and amplify 1/4, 1/4^5^, and 1/4^10^ of the **X**_**20**_ library, respectively, and result in products of reduced diversity. As shown in **Fig. 2g**, using **A**_**20+1**_ caused a right shift of the RT-PCR profile, while **A**_**20+5**_ caused a further shift. The diversity of both PCR products remains high, thus no large difference in end-point signal should be expected. Interestingly, when **A**_**20+10**_ was used, in addition to a further right-shift of the curve, the shape has transformed and resembles that of the **X**_**10**_ library, in good agreement with the reduced diversity of PCR product from 420 (a **X**_**20**_ library) to 410 (a **X**_**10**_ library). When **X**_**10/2**_ was subjected to RT-PCR using **A**_**20+1**_ and **A**_**20+5**_

*Growth with Asymmetric Primers*

While PCR represents a process of population growth, two primers can be treated as essential resources. It is important to note that the two resources are inter-dependent, because limiting one component will damage the exponential growth. This condition of inter-dependent resources has been applied in asymmetric PCR^8^. However, under standard asymmetric PCR conditions, the concentration of one primer is much lower, thus reducing the entire molarity. The kinetics will be governed by the consumption of one primer, preventing the comparison of libraries of different diversity.

We designed a novel asymmetric RT-PCR experiment using primers of various lengths^9^. Shortening primer **A** reduces its annealing temperature to the templates. In the combination of full-length **B’**, the polymerization initiated by the short primer (**A12**, **A15**, or **A18**) will become the rate-as primers, the products’ diversities were reduced from 410 (when the fully complimentary **A20** was used) to 49 and 45, respectively. The difference caused by biased amplification can be observed in both right-shift and shape of the curves (**Fig. 2h**), indicating decreased number of templates as well as diversity. As **A20**, **A**_**20+1**_ and **A**_**20+5**_ are fully complementary to **X**_**10**_, the primers did not affect the amplification curves, unlike **X**_**18**_ (**Extended Data Fig. 5**).limiting step. Using a single DNA as a template, **B’** and **A18** were similarly efficient, thus the combination did not affect the RT-PCR curves remarkably (**Fig. 2i**). However, when **A15** is used, it is still able to initiate chain growth but much less efficiently than **A20**; hence, a dramatically delayed DNA synthesis is observed. When the primer is further shortened to 12 bp (**A12**), the progress is completely abolished. When a highly diverse library is used as template, the simulation experiment predicted that the turning point would be retarded and appear at a lower F value (**Extended Data Fig. 6)**. Because the synthesis of fully complementary duplex becomes slower, the resulting increase of fluorescence is exceeded by the counter effect associated with DNA reshuffling at lower full length DNA concentration. As expected, the RT-PCR course of **A15/B’** illustrates not only the library diversity but also the effect caused by the imbalance between primers (**Fig. 2j**). This design could be used to model conditions in which the inefficient supply of one resource will become decisive for a population’s progress, as in an adaptation scenario^10^.

**Figure 5.**
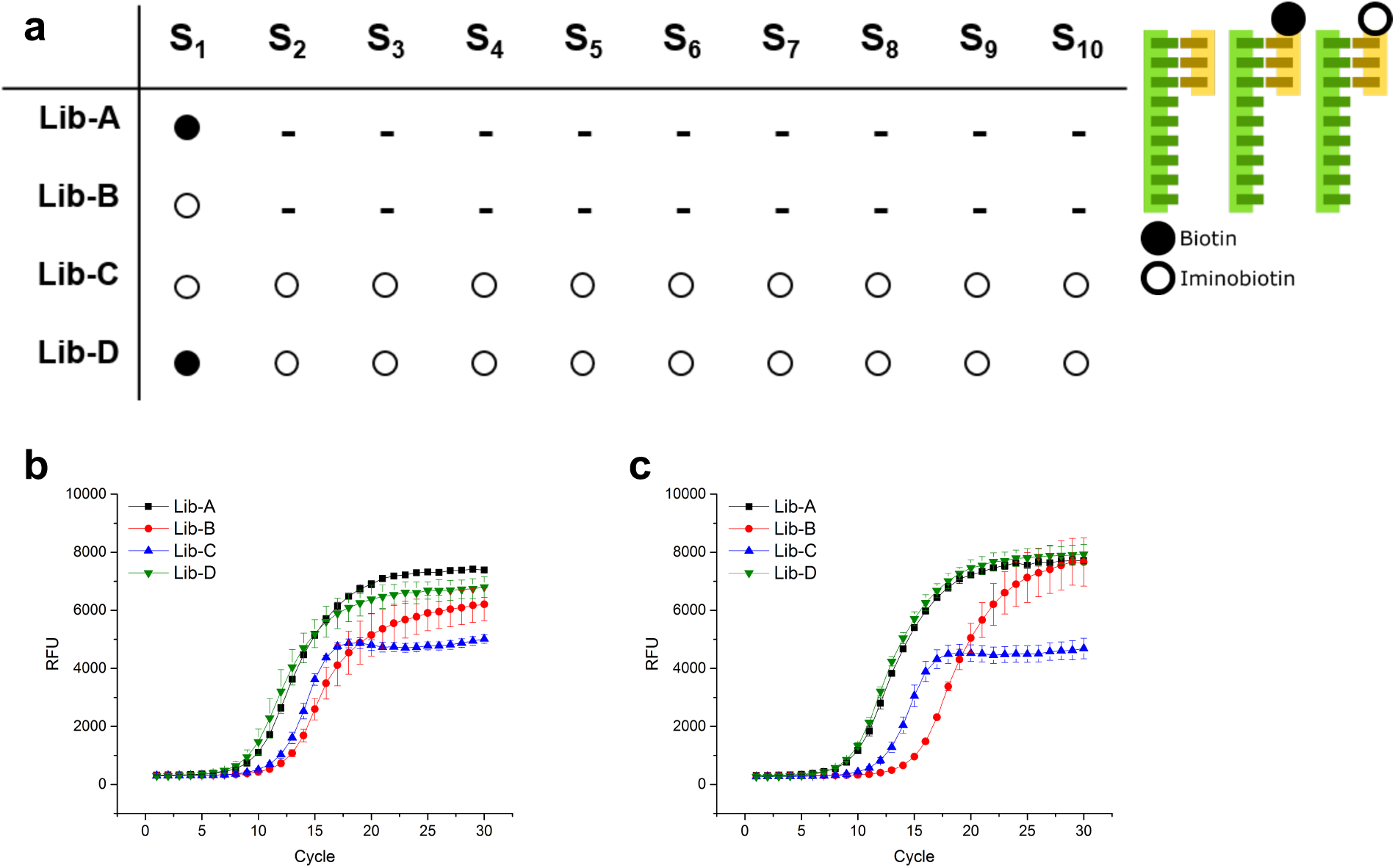
Diversity evaluation of libraries after selection. (**a**) Libraries schematic: ten sequences (**Extended Data Table 1**) were annealed with a 20-mer (**B’**), which was modified with biotin (full dot), iminobiotin (empty dot), or was unmodified. The selected mixture on SA beads after one (**b**) and two washing steps (**c**) was eluted and subjected to RT-PCR for diversity evaluation

### The Protective Effect of High Diversity

To demonstrate that DNA libraries can model not only highly heterogeneous populations under dynamic progress, but also systems subjected to additional selection conditions, we designed the libraries **A-XEcoRV-B** and **A-XSmal-B**. In **XEcoRV** and **XSmaI**, two sequences for the restriction enzymes EcoRV and SmaI were inserted into the randomized regions, respectively (**Fig. 1a**). This provides a mechanism for selection during growth, as the fully complimentary duplexes are the optimal substrates for endonucleases (**Fig.3a**). The mixture of **A-XEcoRV-B** and **A-XSmal-B** was subjected to five or 25 cycles of PCR. Then, a small aliquot of PCR product was treated with one of the restriction enzymes or both. After five cycles, most DNA duplexes were newly synthesized and fully complimentary. Enzyme treatments reduced their concentrations, especially in the presence of both enzymes (**Fig. 3b**). In contrast, after 25 cycles, the effect from enzyme treatment was abolished, as the four curves could not be distinguished from each other (**Fig. 3c**). The emerging complexity caused by DNA reshuffling led to a design of heterogeneous population, which exhibits protective effect against endonuclease digestion (**Extended Data Fig. 7**).

### The Formation of High-Entropic Structures

A mixture of random sequences can cause many potential interactions, and lead to structures other than fully complementary duplex and involving more than two strands. A large library such as **A-X_20_-B** could involve interactions beyond the power of computation. We speculated that in the presence of DNA polymerase a highly diverse DNA library is more likely to evolve complex structures than a simple system^11^. The first indication was an astonishing increase of fluorescence in the RT-PCR course when the sample was pre-treated with 25 PCR cycles (**Fig. 3c**). Without PCR pre-treatment, reoccurrences of fluorescence were observed only after more than 37 RT-PCR cycles. We speculated that whereas the decrease of fluorescence was caused by the generation of less stable mismatching duplexes with low dye binding affinity at elongation temperature, the reoccurrence of fluorescence is caused by the formation of large and complex DNA structures, which are stabilized by the extensive network.

To illustrate the formation of large DNA structures, single DNA sequences or libraries were subjected to 20 or 45 PCR cycles, and the products were analyzed by polyacrylamide gel (PAGE) under denaturing conditions (**Fig. 4a**). After 20 cycles, bands corresponding to DNA larger than 60 bp were observed for the **X**_**16**_ and **X**_**20**_ samples, but not for the sample containing only one sequence (**S1**). After 45 cycles, high molecular weight bands appeared in all samples, though **S1** still produced a strong 60 bp band. Interestingly, the **X**_**16**_ sample produced the largest shift. We then analyzed the PCR products using atomic force microscopy (AFM)^12^. After 20 PCR cycles, mainly small structures of DNA duplex were observed for both **S1** and **X**_**20**_ (**Fig. 4b** and **Extended Data Fig. 8**). Many large structures could be detected after 45 PCR cycles for the samples of **X**_**12**_, **X**_**14**_, **X**_**16**_, and **X**_**20**_, but not that of **S1**. Structures higher than normal DNA duplexes have been observed, especially for the sample using **X**_**16**_ as template. While such structures were not observed with genomic DNA samples^13^, the extensive PCR amplification of DNA libraries of high diversity can evolve to form complex 3D structures.

### Instant Diversity Evaluation of Libraries Under Selection

To demonstrate that DNA libraries can be used not only to build new models, but also as practical tool for existing library technologies, we applied the method to DECL. One major challenge for DECL is to find optimal selection stringency to discover small number of potent binders from large combinatorial libraries, overcoming the low signal-to-noise ratio caused by promiscuous and/or weak interactions between DNA-conjugates and target proteins immobilized on solid support. Unfortunately, a selection experiment will remain as a black box until the hit compounds are revealed after cumbersome steps of decoding and data analysis. We speculated that our diversity analysis method could lead to instant evaluation of library diversity before and after selection.

We have designed four libraries using the ten sequences with the highest diversity (**Fig. 2f**), annealed with a 20-mer (**B’**) modified with biotin (a high affinity binder to the protein streptavidin (SA)) or iminobiotin (a weak binder), or not modified (**Fig. 5a**). **Lib-A** represents a library with a few potent binders without weak binders, and **Lib-B** simulates a library containing a few weak binders. **Lib-C** possesses many weak binders, while **Lib-D** is used to mimic a library with a few high affinity ligands and many weak binders. The libraries were incubated with a SA-coated resin. After removing the supernatant, the resins were subjected to one or two washing steps, representing conditions of different selection stringency.

As shown in **Fig. 5b**, after one washing step, more DNA in **Lib-D** bound to SA-resin than the other libraries, because all DNAs were modified with either strong or weak binders. Interestingly, its end-point is lower than that of **Lib-A**, indicating that the selection using **Lib-D** results in a mixture of compounds, though one compound (**DNA-biotin**) is the major component. While the selection using **Lib-C** enriched more DNA than **Lib-D**, the curve clearly shows that it contains many binders with similar abundance. As expected, the DNA in **Lib-B** is least enriched on SA-resin. The slightly low end-point signal could be caused by the unspecific binding of unmodified DNA.

After the second washing step (**Fig. 5c**), **Lib-A** and **Lib-D** showed that only one construct was bound to the resin. **Lib-C** after selection remained as a mixture of many different compounds. For **Lib-B**, removing unspecific binding of unmodified DNA increased the end-point value to that of **Lib-A**, while the increase of 5.6 amplification cycles is caused by the weak affinity of iminobiotin to SA as compared to biotin. It is important to note that the curves of **Lib-B** and **Lib-D** after one washing step (**Fig. 5b**) could be caused by either two binders of similar abundance or one major binder and many weaker binders in low abundance. Under more stringent selection conditions (two washing steps), the curves of **Lib-B** and **Lib-D** show that they are composed of only one major binder, whereas the curve of **Lib-C**, which is composed of many compounds of similar affinity, remains unchanged.

## CONCLUSION

Degenerated DNA libraries have been widely utilized in biotechnologies such as SELEX and peptide/protein display, while RT-PCR is a standard high throughput method to quantify individual oligonucleotide sequences. The weakened interaction between fluorescence dye and mismatching duplex is also a well-known effect. However, through monitoring the evolving mismatching duplex formation through a PCR course, we can obtain unprecedented insight into the heterogeneity of various DNA libraries.

We designed DNA libraries that behave as synthetic societies and can be used to model heterogeneous populations. The dynamic interactions among the individual members govern the time course and final product of a PCR reaction. Libraries can be designed to simulate different types of diversity and function. For example, heterogeneity in a population has shown a protective effect against enzymatic digestion. The libraries could not only be used to build models to verify hypotheses, but also lead to new discoveries. An unexpected reoccurrence of fluorescence caused by the formation of complex DNA structures has been observed when the libraries were subjected to extended PCR cycles. The resulting amorphous 3D structure produced from a highly diverse library is fundamentally different from the low-entropy structures generated by DNA-origami technology through designing thousands of complementary DNA strands. The RT-PCR method also has applications in biochemical analysis, e.g., to analyze the mixture of oligonucleotides from selection experiment, such as SELEX^14,15^ and DECL^16,17^, as demonstrated in the SA selection experiment (**Fig. 5**).

The application of DNA libraries as synthetic societies will open new venues to model complex and diverse populations. Although it resembles various existing DNA-based computations on the aspect of parallel computing^18^, our method represents a different approach. As compared to some most complex designs of DNA computing using tens to hundreds of unique DNA strands^19^, our library has more than one trillion different sequences and possesses in principle no size limit. It does not aim to perform tasks (in a classical sense) as a calculator. However, it can be used to model systems beyond the power of computation, to simulate the collective behavior of a heterogeneous and dynamic population, and its underlining probabilistic structure associated with an astronomical number of random interactions.

## METHODS

### Reagents and Oligonucleotides

All reagents were purchased from Thermo Fisher Scientific (Germany), unless stated otherwise. All oligonucleotides were purchased from IBA (Germany). Water was nuclease free.

### Annealing

The oligonucleotides were diluted in an annealing buffer solution (10 mM Tris-HCl pH 7.5, 100 mM NaCl, 1 mM EDTA) and heated for 5 min at 95 °C. Then, they were slowly cooled to room temperature. When indicated, the annealed products were incubated with 1X SYBR Green I (Lonza, Switzerland).

### Polymerase Chain Reaction (PCR)

The PCR amplification was carried out in 50 μL in a peqSTAR2x thermocycler (PeqLab, Germany) using the TrueStart(tm) Taq DNA polymerase (2 U) system, with 1.5 mM MgCl2, 150 nM primers, 0.2 mM dNTP mix and 100 pM template. The temperature protocol was: 10 min at 95 °C; 5 or 25 amplification cycles of 15 s at 95 °C, 30 s at 60 °C, and 20 s at 72 °C; 30 s at 72 °C; and 10 s at 20 °C.

### Real-Time PCR (RT-PCR)

The RT-PCR experiments were performed using the PerfeCTa SYBR Green SuperMix, which contains AccuStart(tm)Taq DNA polymerase. The amplification reaction was carried out on a PikoReal(tm) Real-Time PCR System (Thermo Fisher Scientific, Germany), with white 96-well Piko PCR plates and sealed with the respective optical adhesive films. Each reaction well had 10 μL, with 150 nM primers and 100 pM template. The temperature protocol was: 10 min at 95 °C; 20, 30 or 45 amplification cycles of 15 s at 95 °C, 30 s at 60 °C, and 20 s at 72 °C (data acquisition point); 30 s at 72 °C; and 10 s at 20 °C. The results were collected using the PikoReal(tm) Software 2.2.

### Restriction Enzymes

The PCR products were diluted: ten times for the five cycles of amplification sample and 100 times for the 25 cycles of amplification sample. Then, 10 μL of this dilution were incubated with 0.5 μL of a restriction enzyme, SmaI (10 U/μL) and/or an isoschizomer of EcoRV (10 U/μL), for 30 min at 30 °C, 37 min at 30 °C, and 20 min at 80 °C. After, 1 μL of the digested sample was added to the RT-PCR mix (described above) and the amplification was monitored for 30 cycles.

### Denaturing Urea Polyacrylamide Gel Electrophoresis (Urea-PAGE)

A 15% Urea-PAGE (TBE-Urea Gel) was pre-ran in 1X TBE buffer, at 160 V and 10 mA, for 30 min. The RT-PCR samples were added 2X TBE Urea Sample Buffer (final volume of 10 μL) and heated to 70 °C for 3 min; then, immediately placed on ice. The gel ran for 1 h, at 160 V and 10 mA. After the run, the gel was stained for 20 min with 1X SYBR Green II (Lonza, Switzerland) in 1X TBE, and read at 470 nm.

### Atomic Force Microscopy (AFM) Imaging

AFM images of the RT-PCR products were deposited on freshly cleaved mica in the presence of the imaging buffer (300 mM spermidine trihydrochloride, 300 mM NaCl, 20 mM tris(hydroxymethyl)aminomethane in analytical grade water^20^). Optimal dilutions were deposited on the mica (Plano, Germany) and kept still for 90 s. The substrate was then rinsed with analytical grade water and dried with a steady flow of nitrogen^12^. The images were obtained on air, by contact mode, using a Nanowizard II AFM (JPK, Germany). Silicon nitride cantilevers of 200 μm, with a resonant frequency of about 17 kHz and spring constant of about 0.08 N/m with a gold coating were used. Second order polynomial function was used to remove background slope and the images are shown as a heat map of the surface’s topography. DNA origami constructs were imaged as controls to the imaging process.

### Library Selection: Streptavidin

This selection is based on the high affinity binding model for biotin/iminobiotin and SA^21,22^. The DNA libraries (**Fig. 5a** and **Extended Data Table 1**) were annealed to a final concentration of 100 nM. Then, 100 μL of 10 nM DNA libraries were incubated with 10 μL of SA beads (GE Healthcare, UK), at room temperature for 1 h in a shaker, in the selection buffer (150 mM NaCl, 25 mM NaHCO3, 0.005 % Tween 20, pH 9.2). The SA beads were previously washed three times with the selection buffer and the beads slurry was resuspended in selection buffer before being used. After 1 h, the suspension was centrifuged (5 min, 1000 rpm, 4 °C) and the supernatant discarded. Following washing steps, one or two, were performed. The slurry was then subjected to alkaline denaturation (150 nM NaOH, 3 min at room temperature). Immediately after, the suspension was acidified with 1.5 M acetic acid and again centrifuged. One-μL from the resulting supernatant was added to the RT-PCR mix (described above) and the amplification was monitored for 30 cycles.

## ACKNOWLEDGMENTS

This project was supported by the German Bundesministerium für Bildung und Forschung (BMBF) Grant 03Z2E512 and Exist Forschungstransfer (EFT) Grant 03EFFSN091.

## AUTHOR Contributions

H.A. and Y.Z. conceived the project, designed the methods, and experiments; H.A. performed the experiments; Y.Z. wrote the manuscript; A.K.T. performed the AFM imaging; and H.A., W.L. and F.R. designed the libraries. Correspondence and requests for materials should be addressed to Y.Z.

## Extended Data

**Extended Data Figure 1.**
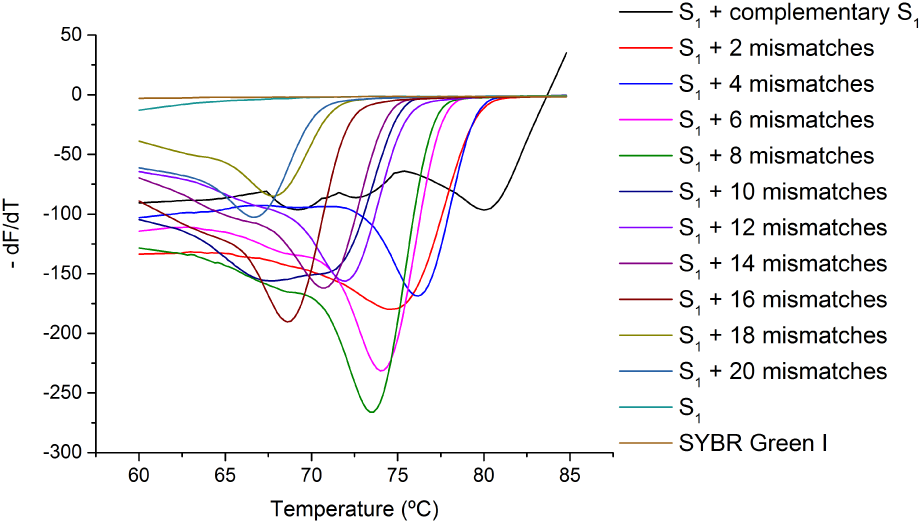
Melting curves for the mismatched libraries.

**Extended Data Figure 2.**
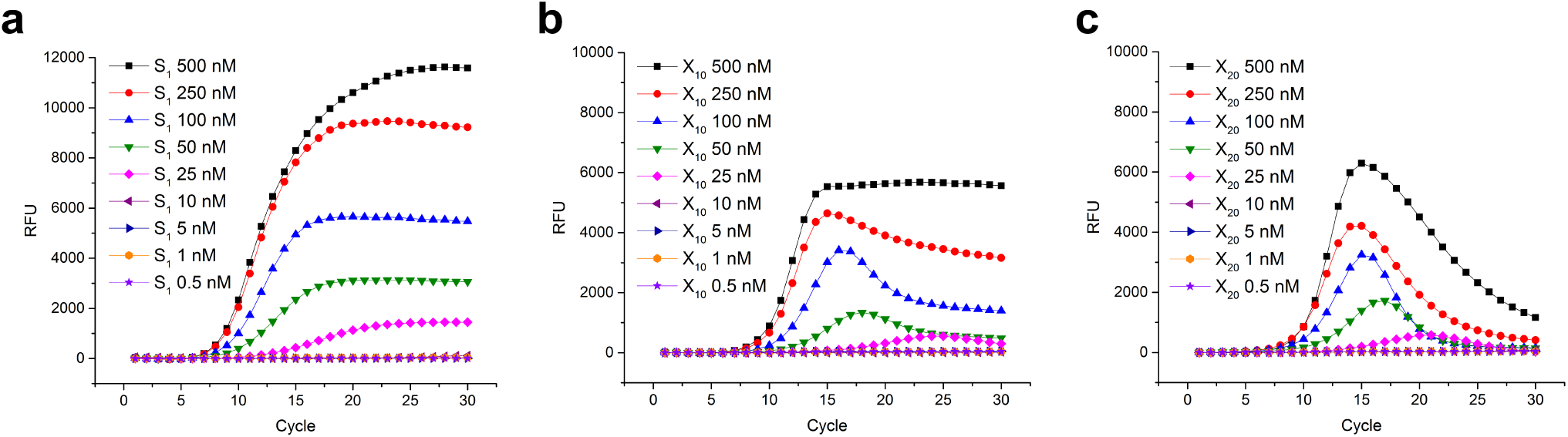
Forward (**A**) and reverse (**B’**) primers titration for **S1** (**a**), **X**_**10**_ (**b**), and **X**_**20**_ (**c**), libraries, with final concentrations ranging from 0.5 to 500 nM for both.

**Extended Data Figure 3.**
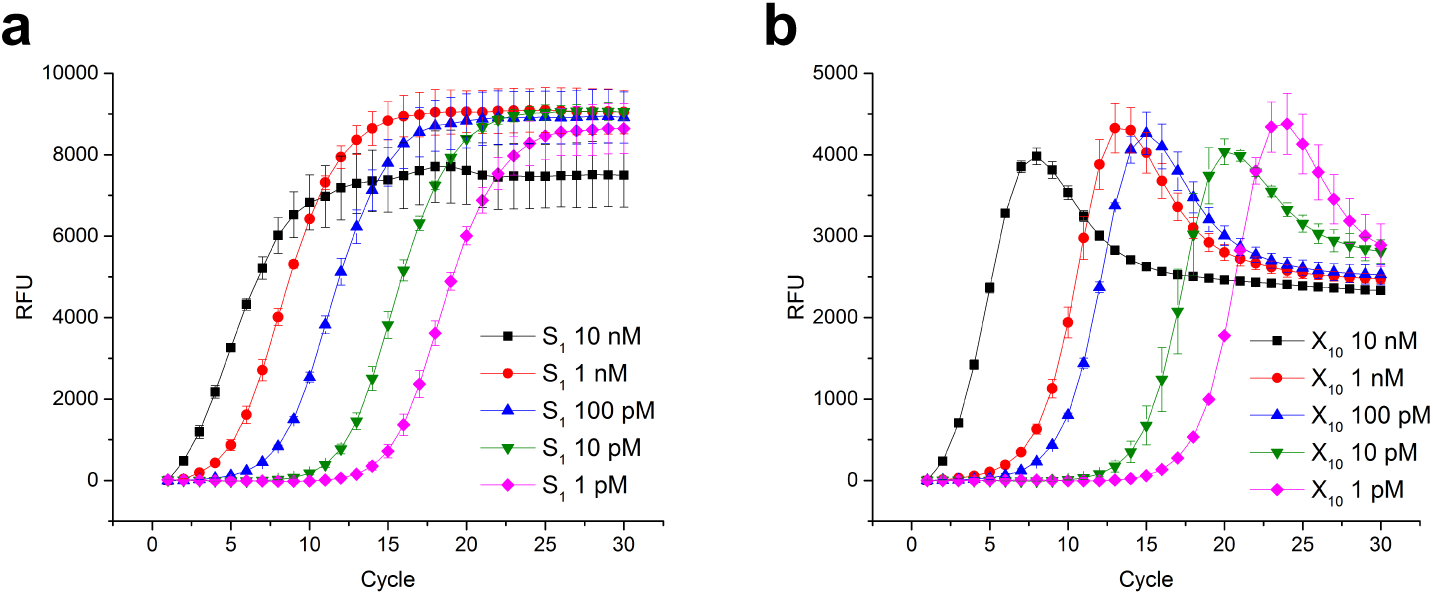
Library titration from 1 pM to 10 nM for **S1** (**a**) and **X**_**10**_ (**b**).

**Extended Data Figure 4.**
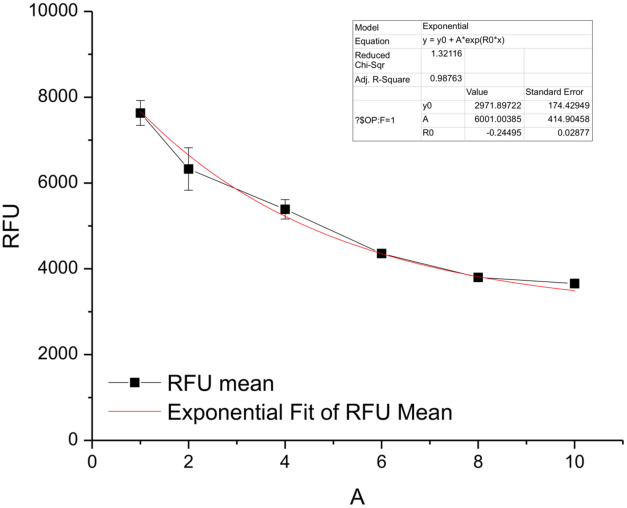
Fluorescence (RFU) at 30 cycles of amplification plotted versus the number of high diversity sequences present in the synthetic mixture. The data was extracted from the graph represented on **Fig. 2f**. The resulting curve was then fitted as an exponential function.

**Extended Data Figure 5.**
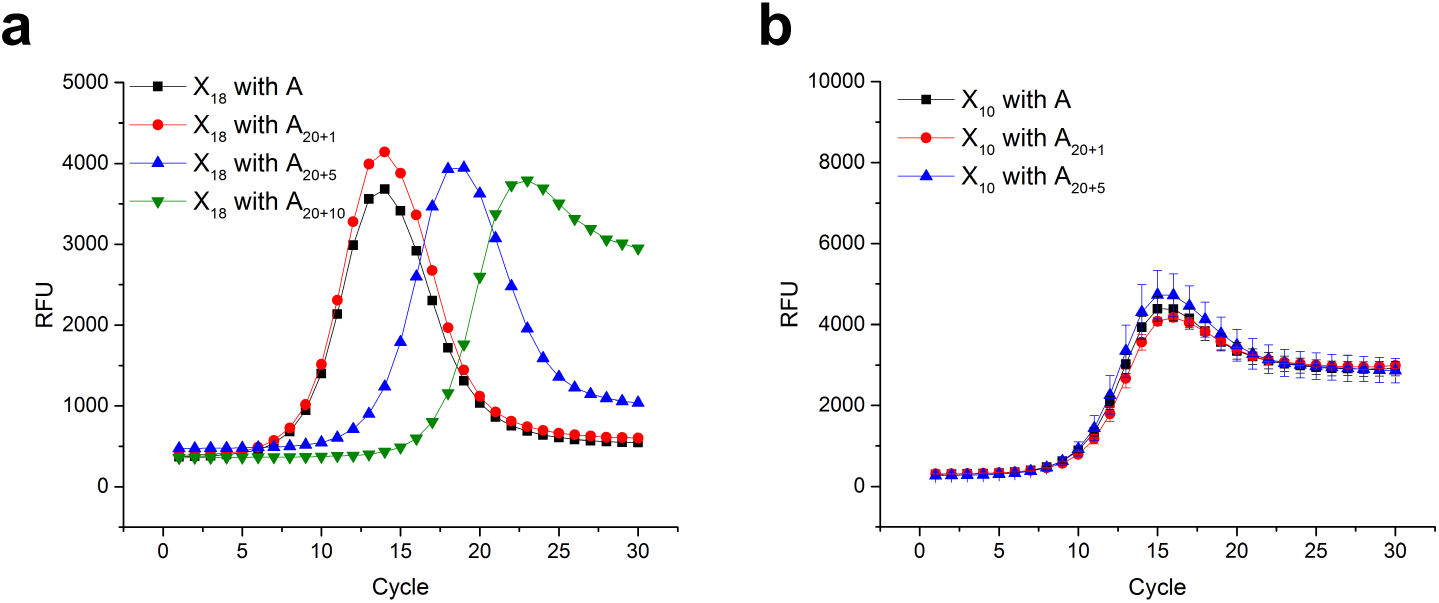
Biased population growth using the longer **A** primers **A**_**20+1**_, **A**_**20+5**_, or **A**_**20+10**_ with the libraries **X**_**18**_ (**a**) and **X**_**10**_ (**b**).

**Extended Data Figure 6.**
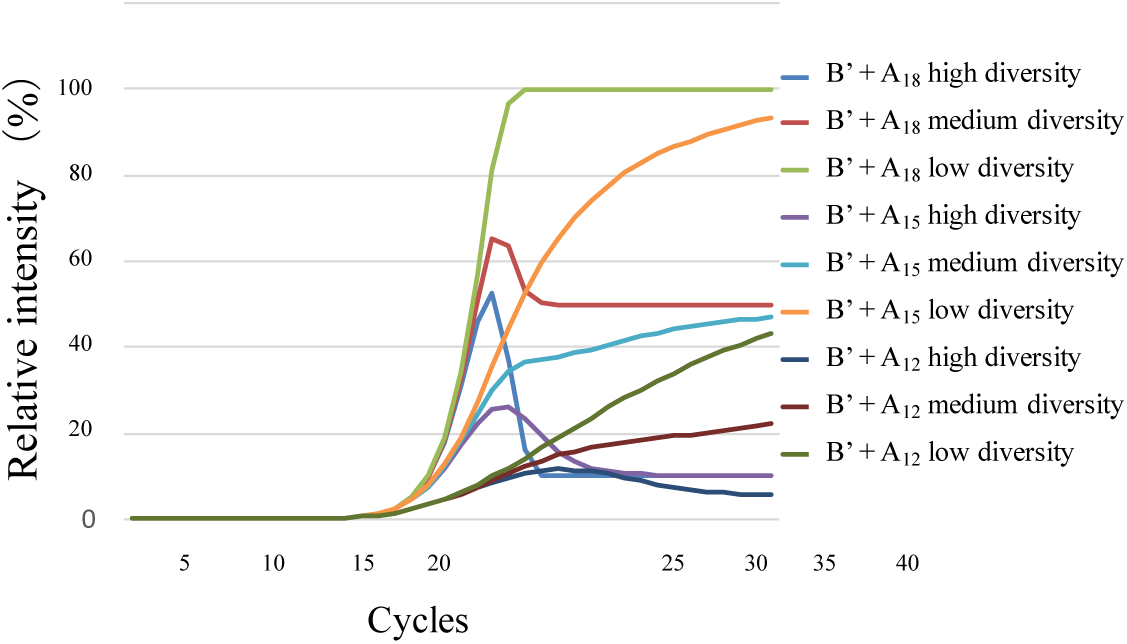
Simulation of PCR course using libraries of different diversity (high, medium and low) and pairs of primer of different lengths (**B’** + **A18**; or **B’** + **A15**; or **B’** + **A12**).

**Extended Data Figure 7.**
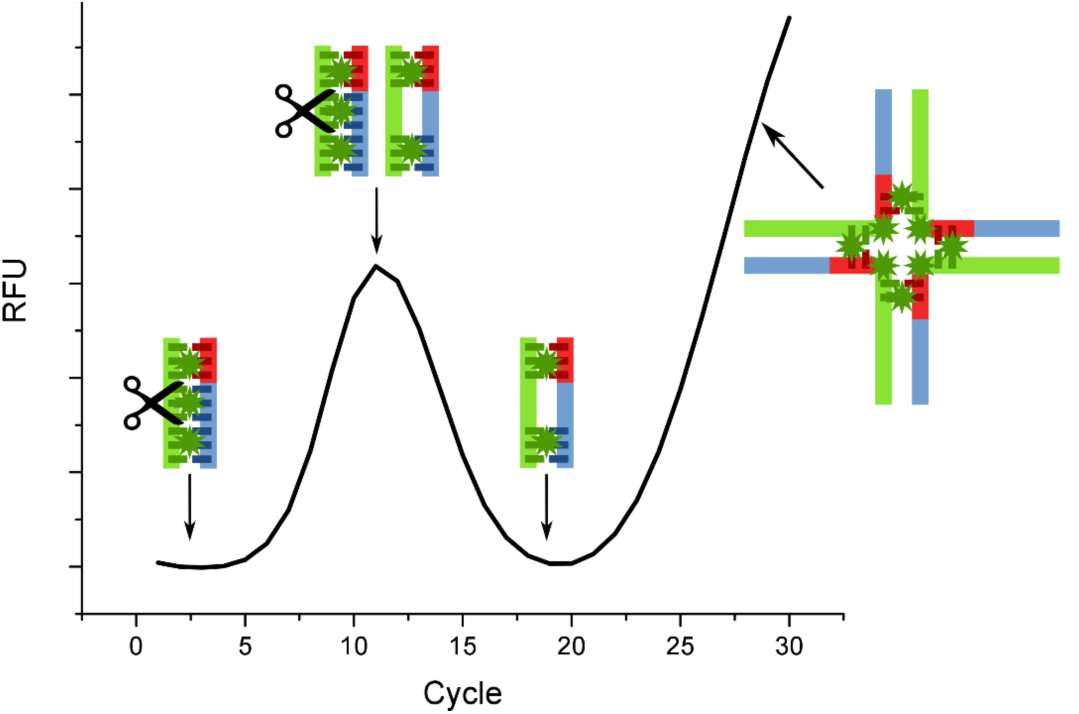
RT-PCR amplification overlook and the impact of diversity on population protection and development of high-entropy structures.

**Extended Data Figure 8.**
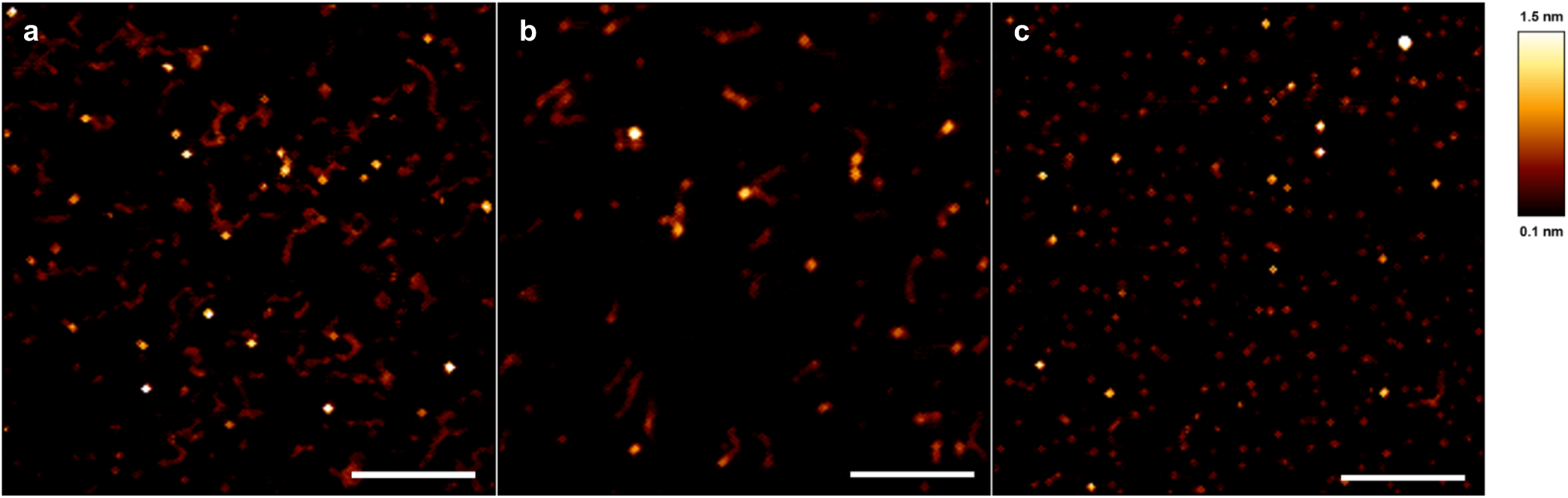
AFM analyses of PCR products of **X**_**12**_ (**a**) and **X**_**14**_ (**b**) after 45 cycles, and **X**_**20**_ after 20 cycles of amplification (**c**). Images are 1μM × 1 μM and the scale bar is 250 nm.

**Extended Data Table 1.**
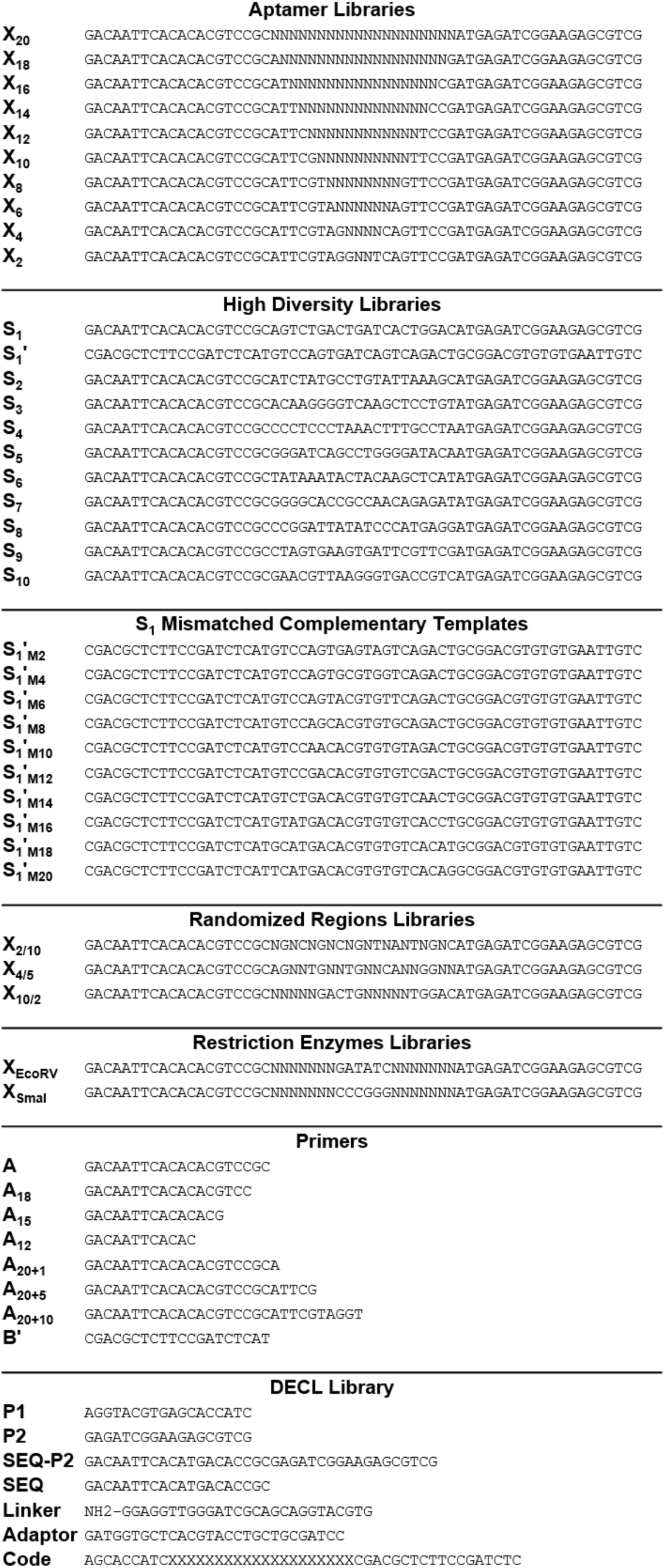
DNA sequences.

